# Comparative genomic analysis of a novel heat-tolerant and euryhaline strain of unicellular marine cyanobacterium *Cyanobacterium* sp. DS4 from a high-temperature lagoon

**DOI:** 10.1101/2025.01.17.633688

**Authors:** Ching-Nen Nathan Chen, Keng-Min Lin, Yu-Chen Lin, Hsin-Ying Chang, Tze Ching Yong, Yi-Fang Chiu, Chih-Horng Kuo, Hsiu-An Chu

## Abstract

**Background:** Cyanobacteria have diversified through their long evolutionary history and occupy a wide range of environments on Earth. To advance our understanding of their adaption mechanisms in extreme environments, we performed stress tolerance characterizations, whole genome sequencing, and comparative genomic analyses of a novel heat-tolerant and euryhaline strain of the unicellular cyanobacterium *Cyanobacterium* sp. Dongsha4 (DS4). This strain was isolated from a lagoon on Dongsha Island in the South China Sea, a habitat with fluctuations in temperature, salinity, light intensity, and nutrient supply.

**Results:** DS4 cells can tolerate long-term high-temperature up to 50 ℃ and salinity from 0 to 6.6 %, which is similar to the results previously obtained for *Cyanobacterium aponinum*. In contrast, most mesophilic cyanobacteria cannot survive under these extreme conditions. Based on the 16S rRNA gene phylogeny, DS4 is most closely related to *Cyanobacterium* sp. NBRC 102756 isolated from Iwojima Island, Japan, and *Cyanobacterium* sp. MCCB114 isolated from Vypeen Island, India. For comparison with strains that have genomic information available, DS4 is most similar to *Cyanobacterium aponinum* strain PCC 10605 (PCC10605), sharing 81.7% of the genomic segments and 92.9% average nucleotide identity (ANI). Gene content comparisons identified multiple distinct features of DS4. Unlike related strains, DS4 possesses the genes necessary for nitrogen fixation. Other notable genes include those involved in photosynthesis, central metabolisms, cyanobacterial starch metabolisms, stress tolerances, and biosynthesis of novel secondary metabolites.

**Conclusions:** These findings promote our understanding of the physiology, ecology, evolution, and stress tolerance mechanisms of cyanobacteria. The information is valuable for future functional studies and biotechnology applications of heat-tolerant and euryhaline marine cyanobacteria.

## Background

Cyanobacteria are great contributors to photosynthesis, biological nitrogen fixation, and the increase of oxygen in the modern atmosphere [1–3]. During their evolutionary history, cyanobacteria of different lineages have occupied and adapted to a wide range of environments on Earth, including hot desert soil crusts, nonacidic hot springs, and arctic/antarctic areas [1, 4]. Some of these cyanobacteria can survive stressful conditions related to light intensity, salinity, temperature, and water availability. However, their adaptation mechanisms to diverse extreme environments are still not fully understood [5]. These cyanobacteria and their natural bioproducts, including carotenoids, phycobilins and lantipeptides (lantibiotics), have been utilized in numerous biotechnology applications [6, 7]. Additionally, saltwater cyanobacteria hold significant potential for biotechnological and bioremediation applications, especially as freshwater resources become increasingly scarce due to the impacts of climate change [5, 6].

Dongsha Atoll is a coral reef atoll located in the South China Sea [8]. Dongsha Island, situated on the western part, is the only above-water part of the atoll during high tide. This island has a small lagoon (at 20° 70’N and 116° 80’E) with an average depth of about one meter. The surface seawater temperature around Dongsha Atoll in a year is between 22 and 32 °C [9]. However, due to the effects of strong sunlight and poor water exchange, the seawater temperature in this lagoon is higher than 35 °C constantly during daytime and can rise over 38 °C in summer [9]. This unusually high seawater temperature makes this location a ground of natural selection for thermotolerance [10]. A new species of thermotolerant microalga, *Tetraselmis* sp. DS3, was isolated and characterized recently from this location [10]. The DS3 cells could grow at 40 °C, the highest temperature reported for marine microalgal growth. In addition, this small lagoon is subject to seasonal changes and daily fluctuations in temperature, salinity, light and nutrient supply. Therefore, a thermotolerant cyanobacterium isolated from this site may be also tolerant to other kinds of abiotic stresses. The new isolate might be an attractive biological system for cellular and biochemical study of these tolerance mechanisms, as well as for potential biotechnology applications.

In this work, we report the isolation and stress tolerance properties of a novel heat-tolerant, euryhaline, unicellular cyanobacterial strain Dongsha4 (DS4) from the lagoon of Dongsha Island. Phylogeny analysis based on 16S rRNA genes showed that DS4 is likely a member of the genus *Cyanobacterium* [11, 12]. This new isolate could grow in media containing 3% salt at 50 °C. This stress tolerance is unusual among the reported mesophilic cyanobacteria but shared with the closely related thermotolerant species *Cyanobacterium aponinum*. To investigate the distinctive genetic features and underlying adaptation mechanisms of this new strain, we sequenced the whole genome and conducted comparative genome analysis of DS4 with related cyanobacteria that have genome sequence information available.

## Methods

### Isolation of DS4 from Dongsha lagoon

Seawater samples were collected from the small lagoon of Dongsha Atoll (20°70’N, 116°72’E) in the South China Sea. Cells in the samples were collected using centrifugation at 3000 x *g* for 10 min at room temperature, and spread onto 1.5% agar plates. These plates were prepared with modified f medium in which the levels of nitrate and phosphate were changed to 3.5 mM NaNO_3_ and 0.15 mM NaH_2_PO_4_, respectively, and Na_2_SiO_3_ was dropped [13]. These plates were incubated under continuous 80 μmol photons m^-2^s^-1^ white light at 28 °C for five days, and then 32 °C under the same illumination conditions to grow single colonies. The DS4 strain was isolated and chosen for this study.

### Growth conditions

The DS4 strain was grown axenically at different temperatures, in BG11 or modified BG11 (mBG11, with 3.3% NaCl) liquid medium [14] according to the exact experiments as specified in the Results section. These cultures were bubbled with filtered air (pore size 0.2 μm, Millipore) in an orbital shaker (speed 150 rpm). The light intensity was 30 μmol photons m^-2^s^-1^.

### Microscopy

The procedures for Scanning Electron Microscopy (SEM) and Transmission Electron Microscopy (TEM) were described previously [15].

### DNA extraction

Total genomic DNA was isolated from 150 mL of DS4 culture grown for 4-5 days. The DNeasy Plant Maxi Kit (QIAGEN, Germany) was used to extract the DNA that was used for Illumina MiSeq sequencing. To obtain high quality DNA for Oxford Nanopore Technologies (ONT) MinION sequencing, a modified CTAB method and further purified with DNeasy Blood & Tissue Kit (QIAGEN, Venlo, Netherland) was used as described in Chen et al. [16]. Quality and quantity of the purified genomic DNA samples were assessed by using the NanoDrop 2000 spectrophotometer (ThermoFisher, USA) and 1% agarose gel electrophoresis.

### Genome sequencing

The procedures for genome sequencing and analysis were based on those described in our previous work [15, 18, 19]. All bioinformatics tools were used with the default settings unless stated otherwise. Further details of the methodology are provided below.

The raw reads from both platforms were combined for *de novo* assembly by using Unicycler v0.4.9b [20]. For validation, the Illumina and ONT raw reads were mapped to the assembly by using BWA v0.7.12 [21] and Minimap2 v2.15 [22], respectively. The results were programmatically checked by using SAMtools v1.2 [23] and manually inspected by using IGV v2.3.57 [24]. The finalized assembly was submitted to the National Center for Biotechnology Information (NCBI) and annotated by using their Prokaryotic Genome Annotation Pipeline (PGAP) v5.3 [25].

### Comparative analysis

For comparative analysis, the 16S rRNA gene of DS4 was used as the query to run BLASTN [26] against the NCBI Genbank 16S rRNA gene sequences and nucleotide collection (nt) databases [27]. Related strains with genome sequences available were identified (Table 1). To quantify genome similarity, FastANI v1.1 [28] was used to calculate the percentage of genomic segments mapped and the average nucleotide identity (ANI) based on these segments for each genome pair; only the chromosomes were included in this analysis and the plasmids were excluded. The homologous gene clusters (HGCs) among these genomes were inferred using OrthoMCL v1.3 [29]. For pairwise genome alignment, the NUCleotide MUMmer (NUCmer) program of the MUMmer package v3.23 [30] was used with the setting “–maxmatch –mincluster 2000.”. Additionally, MAUVE v2015-02-25 [31] was also used for chromosome level alignment.

**Table 1.**
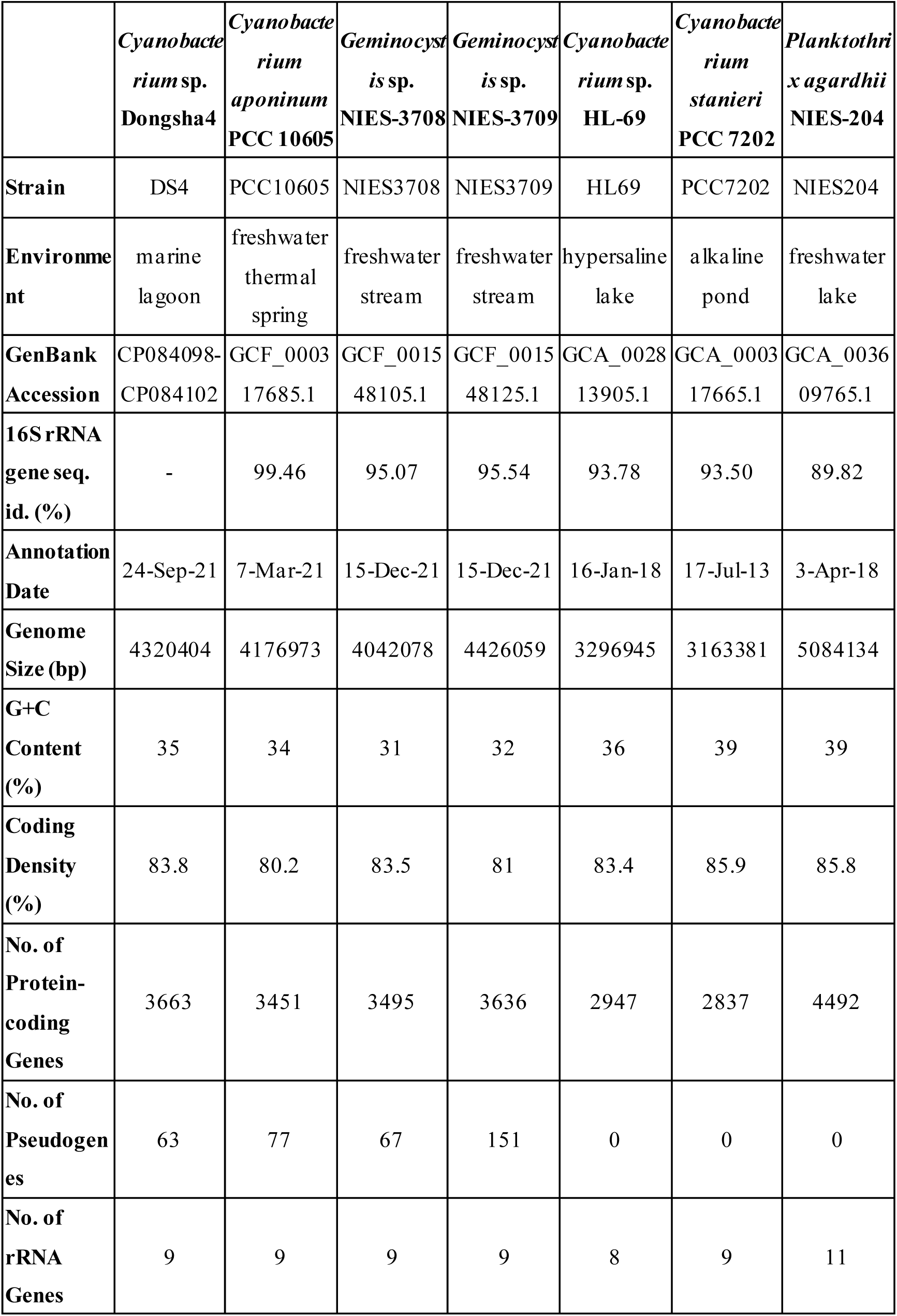
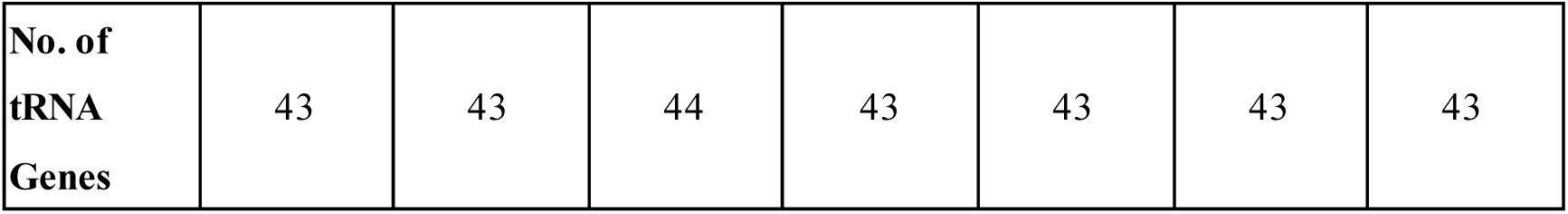
Genome statistics of representative closely related cyanobacteria. The 16S rRNA gene sequence identity values were calculated based on comparisons with DS4.

For phylogenetic inference, the multiple sequence alignment was conducted using MUSCLE v3.8.31 [32]. The maximum likelihood inference was conducted using PhyML v3.3 [33]. The proportion of invariable sites and the gamma distribution parameter were estimated using the dataset, and the number of substitute rate categories was set to four.

## Results and discussion

### Stress tolerance properties

The strain DS4 exhibited distinct heat tolerance properties compared with other mesophilic cyanobacteria. The DS4 cells showed normal photosynthetic growth at 30 °C under both BG11 and mBG11 (with 3.3 % NaCl) growth conditions and maintained apparent photosynthetic growth under mBG11 growth conditions up to 50 ℃ (Figure 1A). In addition, 3 hours of 50 °C thermal stress followed by 21 hours of 35 °C recovery each day did not slow down the growth rate of the DS4 cells. Moreover, DS4 cells can tolerate high temperature stress at 70 ℃ for 3 hrs and the mortality rate of the cells was only 7.1 ± 1.8% (n = 3) (Figure 1 and Supplementary Figure 1). In comparison, previous studies reported that *Cyanobacterium aponinum* PCC 10605 and other related strains can grow at temperature up to 45-50 ℃ [10, 34]. PCC 10605 was isolated from Euganean thermal springs in Italy and is the type strain for *C. aponinum* [12]. *C. aponinum* is a thermotolerant cyanobacterial species found in different thermal springs and desert environments [12, 34–38]. Because of their thermotolerant properties, *C. aponinum* strains have been widely used for scientific researches, including biotechnological applications [35–38]. In contrast, the model mesophilic cyanobacterium *Synechocystis* sp. PCC6803 cannot grow above 43 ℃ [39, 40] and can only tolerate short-term heat stress up to 50 ℃ [39, 40].

**Figure 1.**
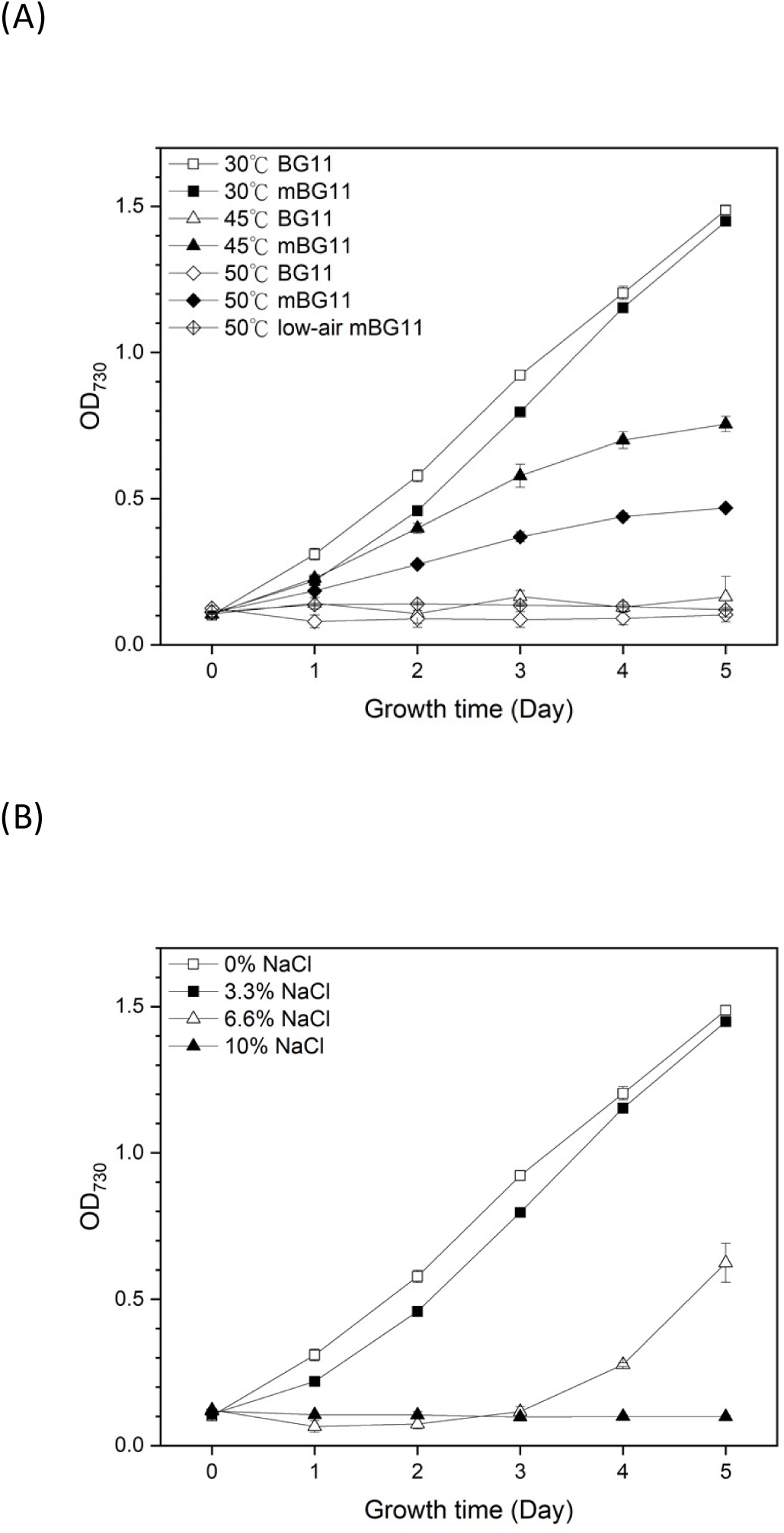
Stress tolerance properties of DS4 cells. (A) Photosynthetic growth of Cyanobacterium sp. DS4 at different temperatures (30, 45 and 50 °C) and different salinity [in BG11 or mBG11 (with 3.3% NaCl) liquid medium]; (B) Photosynthetic growth of DS4 cells in BG11 liquid medium supplemented with different levels of NaCl at 30 °C. All cultures were bubbled with filtered air in an orbital shaker (speed 150 rpm), except for “low-air mBG11” cultures in panel (A), which were not bubbled with air. The light intensity was 30 μmol photons m-2s-1. Data are the mean ± SD (n = 2).

Furthermore, DS4 cells were able to grow at 30 ℃ in mBG11 medium with salt levels from 0 to 6.6% NaCl (Figure 1B). Thus, DS4 showed euryhaline (halo- and osmotic tolerant) properties. Moreover, the growth of DS4 cells at 50 ℃ required the saline growth medium (*e.g.*, 3.3% NaCl or sea salts) and bubbling with air (Figure 1A). Salinity therefore seemed to have a positive effect on thermotolerance of DS4. Similar effects of salinity on thermotolerance have been reported in some hot-spring cyanobacteria, including *C. aponinum* [34, 41]. These stress tolerant properties may help DS4 adapt and survive under harsh environmental conditions, such as fluctuations in temperature, salinity, and other environmental factors, in the small lagoon of Dongsha Island.

### Cell morphology

The images of SEM, TEM, and light microscopy showed that the DS4 cells were typically round to ellipsoid in shape, about 2-3 μm long (Figure 2 and Supplementary Figure 1). DS4 cells at the stage of binary fission were observed. Especially at 45 ℃ and in BG11 liquid medium, the shapes of many DS4 cells were elongated. This is possibly due to impairments in cell fission (Figure 2C and G). In addition, the SEM images showed that the surface of DS4 cells was covered with a thick cell wall, presumably an extracellular polysaccharide layer (Figure 2A-D). The TEM images also showed the irregular thylakoid structures in the DS4 cells. Fimbriae-like protrusions are present on the surface of DS4 (Figure 2E-H).

**Figure 2.**
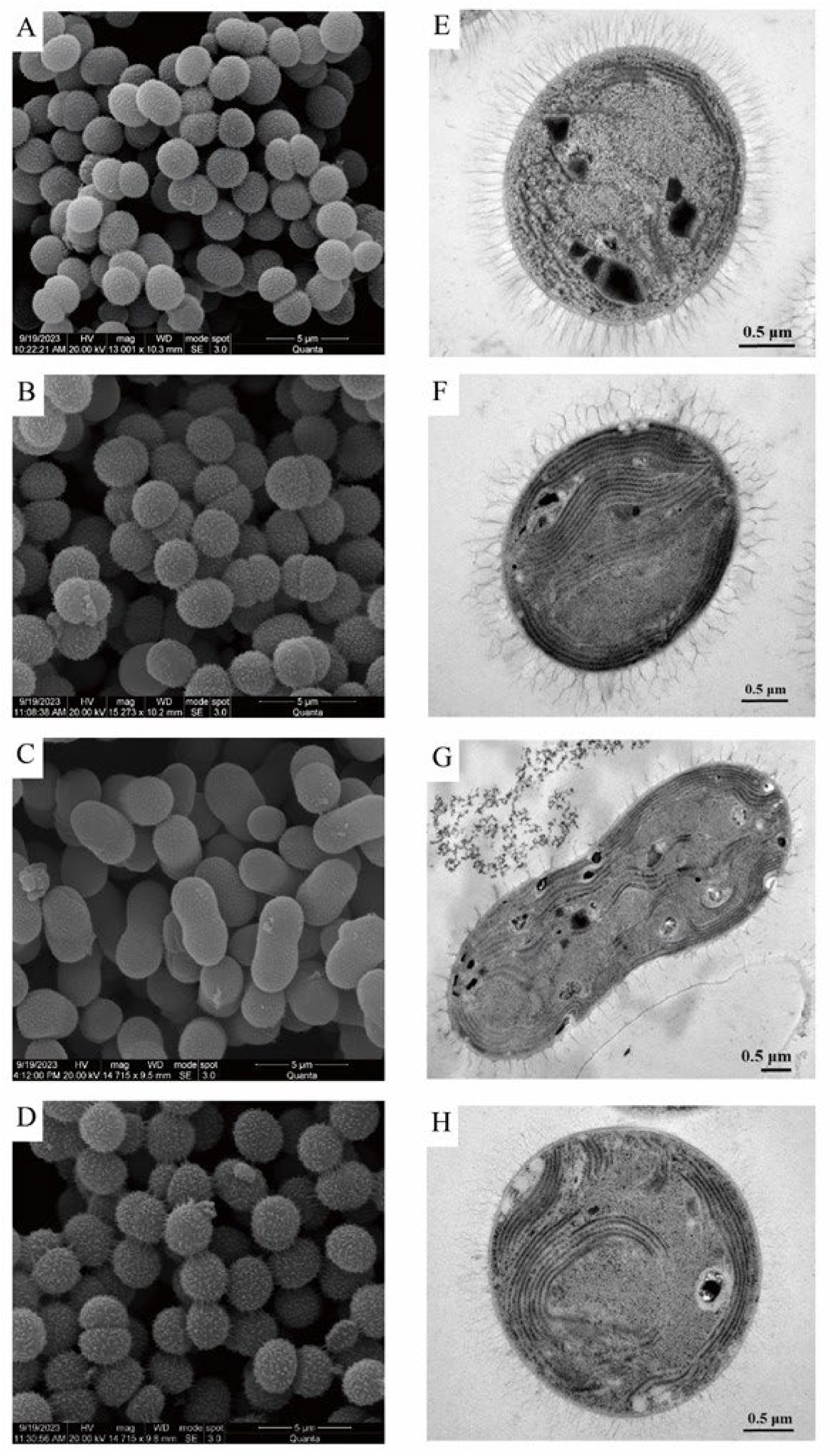
Scanning electron micrograph (SEM) and transmission electron micrograph (TEM) of *Cyanobacterium* sp. DS4 cells grown under different temperatures and different salinity (BG11 and mBG11) conditions. All cultures were bubbled with filtered air in an orbital shaker (speed 150 rpm). (A-D) SEM images; (E-H) TEM images. (A) and (E), at 30 °C in the BG11 liquid medium; (B) and (F), at 30 °C in the mBG11 liquid medium; (C) and (G), at 45 °C in the BG11 liquid medium; (D) and (H), at 45 °C in the mBG11 liquid medium.

### Genome sequence

The whole-genome shotgun sequencing generated 1,865,196*2 pair-end Illumina reads and 43,869 ONT long reads (average length = 28,001 bp, N50 length = 38,678 bp, total length = 1,228,373,655 bp). These reads provided ∼260- and ∼284-fold coverage based on the Illumina and ONT data set, respectively. The complete genome sequence of DS4 contains one circular chromosome that is 4,250,085 bp in size and has 34.8% G+C content. Additionally, four circular plasmids were found, which are 26,702, 20,643, 13,205, and 9,769 bp in size, respectively. For gene content, the annotation includes 43 tRNA genes, three complete set of 16S-23S-5S rRNA genes, and 3,663 intact protein-coding genes. These genomic properties are similar to those found in other related strains, particularly *Cyanobacterium aponinum* PCC 10605 (Table 1).

### Phylogenetic placement

Based on the 16S rRNA gene phylogeny (Figure 3A), DS4 is most closely related to *Cyanobacterium* sp. NBRC 102756 (NBRC 102756, previously named as MBIC 10216 or *Synechocystis aquatilis* SI-2) isolated from Iwojima Island, Japan [11, 42] and to a few strains isolated from Vypeen Island, India (Cyanobacterium sp. MCCB114/115/238) [43, 44]. NBRC 102756 is known to be able to accumulate amylopectin-like polysaccharides in cells, designated as cyanobacterial starch [11]. However, the genome information of NBRC 102756 and MCCB114 strains is unavailable. Among the cyanobacteria with genome sequences available, the DS4 has the highest sequence similarity with *Cyanobacterium aponinum* PCC 10605.

**Figure 3.**
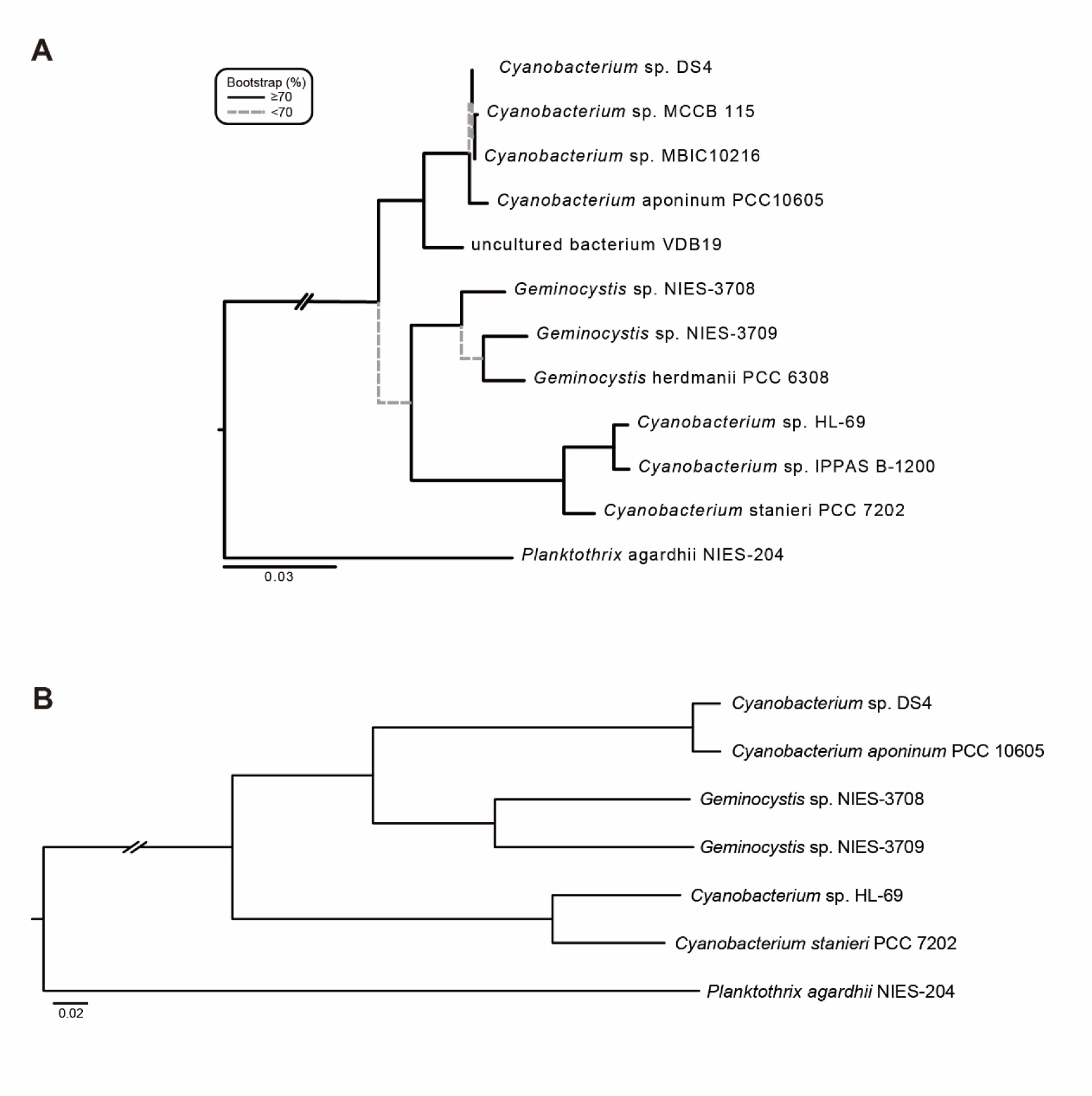
Maximum likelihood phylogeny containing DS4 and related strains. Levels of bootstrap supports were estimated based on 1,000 replicates. (A) 16S rRNA gene phylogeny. Branches with bootstrap support lower than 70% are plotted with grey dashed lines. (B) Core genome phylogeny. All branches received 100% bootstrap support.

### Comparisons with other strains

Based on the 16S rRNA gene phylogeny (Figure 3A), we selected the related strains that have genome sequences available for comparative analysis (Table 1). These include three strains classified as *Cyanobacterium* [45] and two strains classified as *Geminocystis* [46, 47]. In addition to these closely related strains in the order Chroococcales, *Planktothrix agardhii* NIES-204 from the order Oscillatoriales was included as the outgroup [48].

For overall genome similarity, *C. aponinum* PCC 10605 is the most closely related strain of the DS4, sharing 81.7% of the genomic segments and 92.9% ANI (Figure 4). This ANI value is below the 95% threshold suggested for delineation of bacterial species [28], indicating that DS4 may not be a member of *C. aponinum*. However, to reach a more solid conclusion regarding species assignment, comparisons with additional strains within this genus upon genome sequence availability is needed.

**Figure 4.**
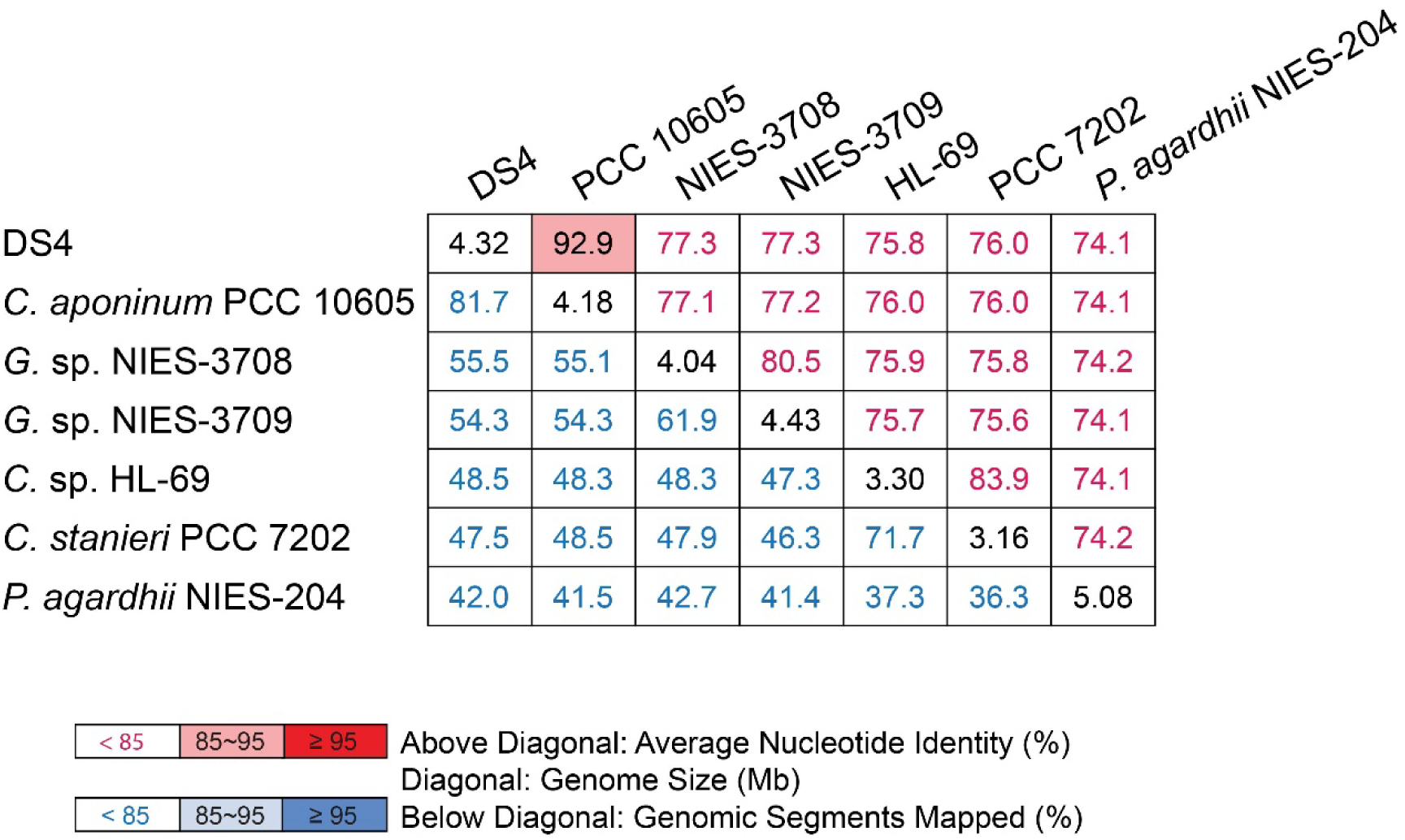
Average nucleotide identity (ANI) analysis of DS4 and related cyanobacterial strains with complete genome assemblies available. For the pairwise comparisons, the ANI value (%) and the proportion of genomic segments mapped (%) are indicated in the cells above and below the diagonal, respectively. The genome sizes (Mb) are indicated in the cells along the diagonal.

The genome alignment between DS4 and PCC 10605 revealed no notable synteny conservation (Figure 5). Such lack of synteny conservation in chromosome organization has been reported previously for other cyanobacteria [15, 48, 49].

**Figure 5.**
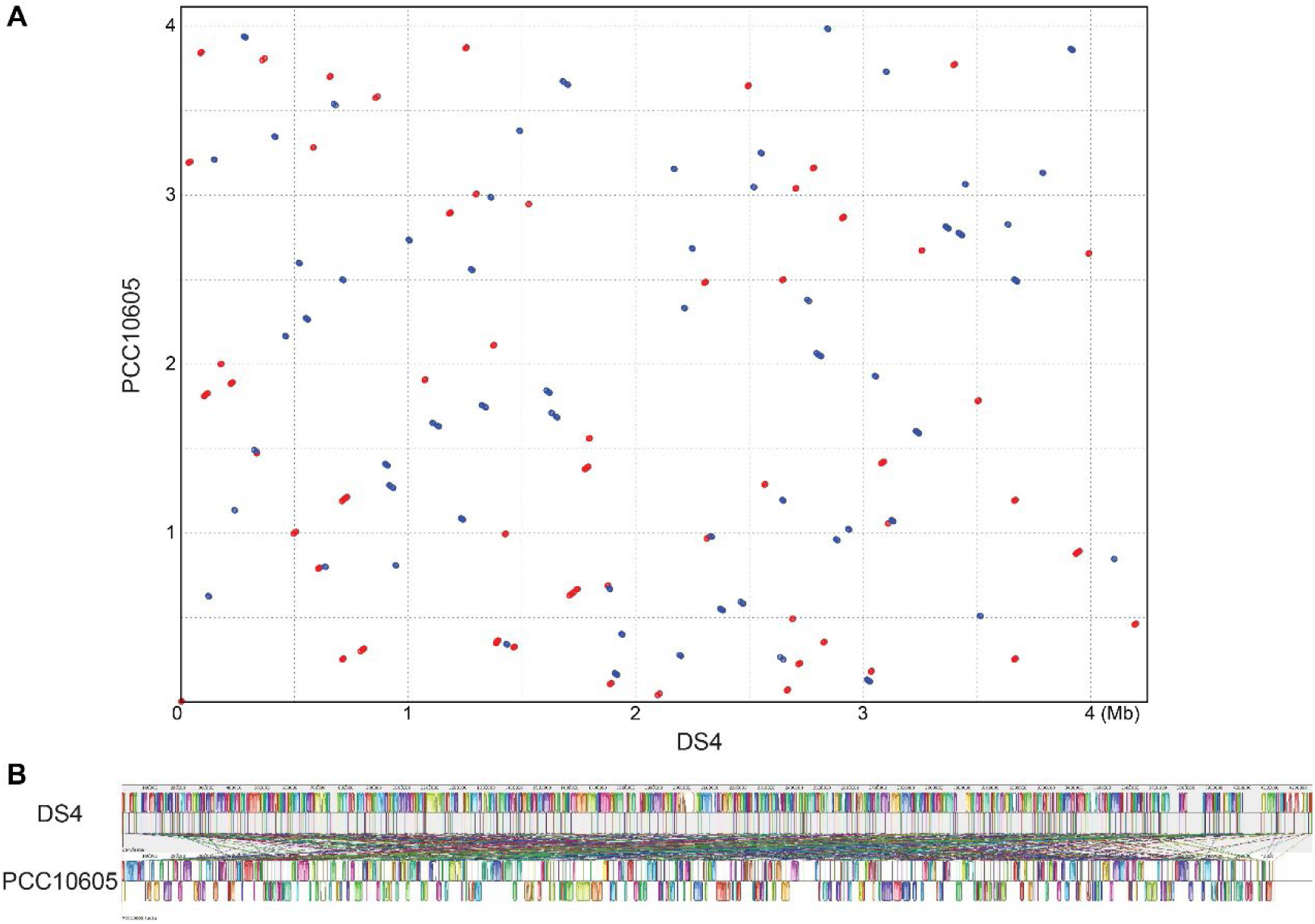
Genome alignments between DS4 and PCC 10605. (A) Alignment inferred by MUMmer. Matched regions with the same and opposite orientations are indicated by red and blue dots, respectively. (B) Alignment inferred by MAUVE. The strain DS4 is used as the reference. Matched regions are indicated by the same color in both assemblies and linked by a line connecting the two regions. In PCC 10605, matched regions with the same and opposition orientations with DS4 are plotted above and below the black line in the center, respectively.

For gene content comparison, 1,610 homologous gene clusters (HGCs) were shared by all of the strains compared. Among these HGCs, 1,525 were conserved as single-copy genes in all strains. The concatenated alignment of these 1,525 HGCs contained 518,506 aligned amino acid sites and was used to infer a maximum-likelihood phylogeny that received 100% bootstrap support in branches (Figure 3B). Consistent with the results from the 16S rRNA gene phylogeny and the ANI analysis, the DS4 is most closely related to PCC 10605. The two *Geminocystis* strains NIES-3708 and NIES-3709 collected in Japan form the sister group, while the other two *Cyanobacterium* strains PCC 7202 and HL-69 collected in the USA appeared to be more distantly related. In addition to the phylogenetic divergence, the two *Cyanobacterium* strains collected in the USA both have much smaller genome sizes (Figure 4). These findings, together with the 16S rRNA gene phylogeny, indicated that the genus *Cyanobacterium* is a polyphyletic group that contains diverse lineages, and future re-classification of strains or taxonomic revisions is required to avoid confusion [50].

### Overview of the notable genetic differentiations among DS4 and other strains

Notable genetic differentiations among the strains compared are summarized in Figure 6. The strain DS4 has many unique genomic features which are different from those in the genome of PCC 10605 and other closely related strains, including gene clusters involved in nitrogen fixation, photo-protection, and novel natural product biosynthesis. In addition, DS4 and PCC10605 shared many distinctive genomic features, including many distinct genes/gene clusters involved in cyanobacterial starch metabolism, central metabolism, hydrogen metabolism, thermotolerance, salt tolerance as well as lantipeptide/bactericide biosynthesis and export systems. More detailed information regarding the inferred function, distribution patterns, and molecular evolution of these genes are provided in the following sections.

**Figure 6.**
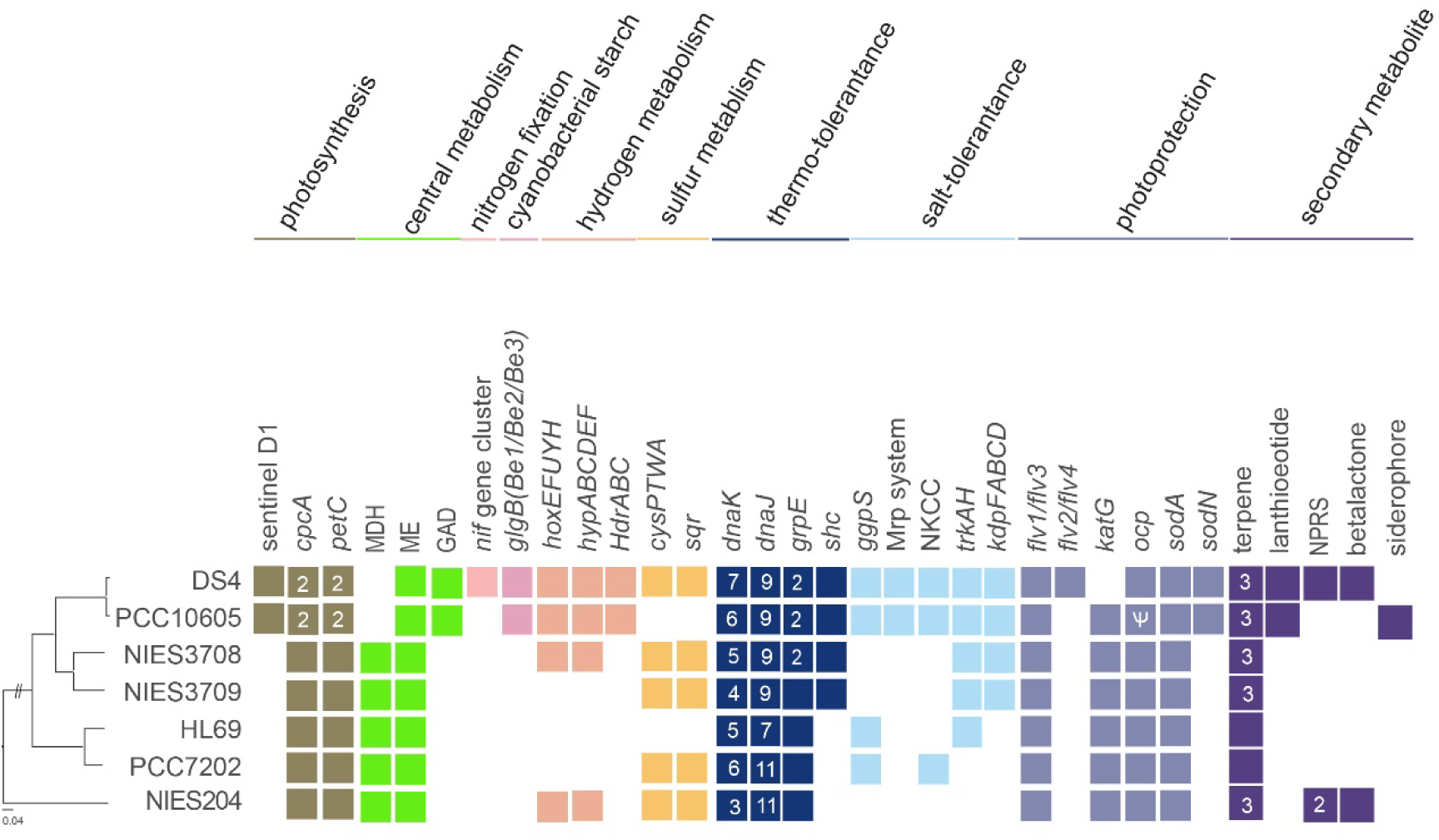
Overview of notable gene content. Gene presence and absence are indicated by colored and white background, respectively. Multi-copy genes are indicated by a dark-colored background with the copy numbers labeled.

### Nitrogen fixation

One of major distinct features in the DS4 genome is the presence of a large gene cluster for nitrogen fixation (Dongsha4_16200-16345) (Figure 6). This nitrogen-fixing (*nif*) gene cluster contained 29 predicted genes which included *nifBSUHDKEN* encoded for a Mo dependent nitrogenase system. This *nif* gene cluster was not found in PCC10605 and several other closely related strains (Figure 6). The amino acid sequence of NifH and NifDK in DS4 are identical to those in *Cyanobacterium.* sp. NBRC10276 and show about 90% amino acid sequence identity to NifH and NifDK genes in baeocystous cyanobacteria, such as *Xenococcus* sp. PCC 7305 and *Chroococcidiopsis* sp. PCC 6712 (Supplementary Figure 3) [51].

### Photosynthesis

The major structural components of photosynthesis genes are generally conserved in the DS4 genome. DS4 has phycobilisome genes for the allophycocyanin core and phycocyanin subunits, but not for phycoerythrin subunits. One of the distinctive features in DS4 (also in PCC10605) is the presence of the atypical *psbA* gene (Dongsha4_01220) (Figure 6) which encoded a predicted sentinel D1 protein with several mutations on essential manganese ligands to the oxygen-evolution complex (Supplementary Figure 2) [52, 53]. The expression of sentinel D1 protein during the dark period may protect and enable nitrogen fixation and hydrogenation reactions which are highly oxygen sensitive processes. In addition, DS4 and PCC10605 havetwo copies of cytochrome *b*6-F complex Rieske iron sulfur protein (*petC*) genes, whereas most cyanobacterial genomes have only one copy.

### Central carbon metabolism

The cyanobacterial tricarboxylic acid (TCA) cycle does not use the typical α-ketoglutarate dehydrogenase but rather uses α-ketoglutarate decarboxylase (KGD), which sequentially converts α-ketoglutarate into succinic semialdehyde and succinate [54]. DS4 and PCC10605 have one distinct gene for glutamate decarboxylase (GAD) which is absent in most cyanobacterial genomes (Figure 6). The GAD enzyme converts glutamate to γ-aminobutyric acid (GABA) [55]. Thus, DS4 and PCC10605 may be able to use the GABA shunt pathway to complete the TCA cycle independent of the KGD bypass (Figure 7). In addition, DS4 and PCC10605 genomes do not have the gene for malate dehydrogenase (MDH), which catalyzes the NAD/NADH-dependent interconversion of malate and oxaloacetate. A recent study reported that malic enzyme (ME), not MDH, is mainly responsible for the oxidization of malate from the TCA cycle in cyanobacteria (Figure 7) [56]. Thus, MDH may be dispensable and may have been lost during the evolution of DS4 and PCC10605 genomes. It would be interesting for future research to determine whether the GABA shunt and the absence of MDH may contribute to the stress tolerance mechanisms of DS4 and PCC10605.

**Figure 7.**
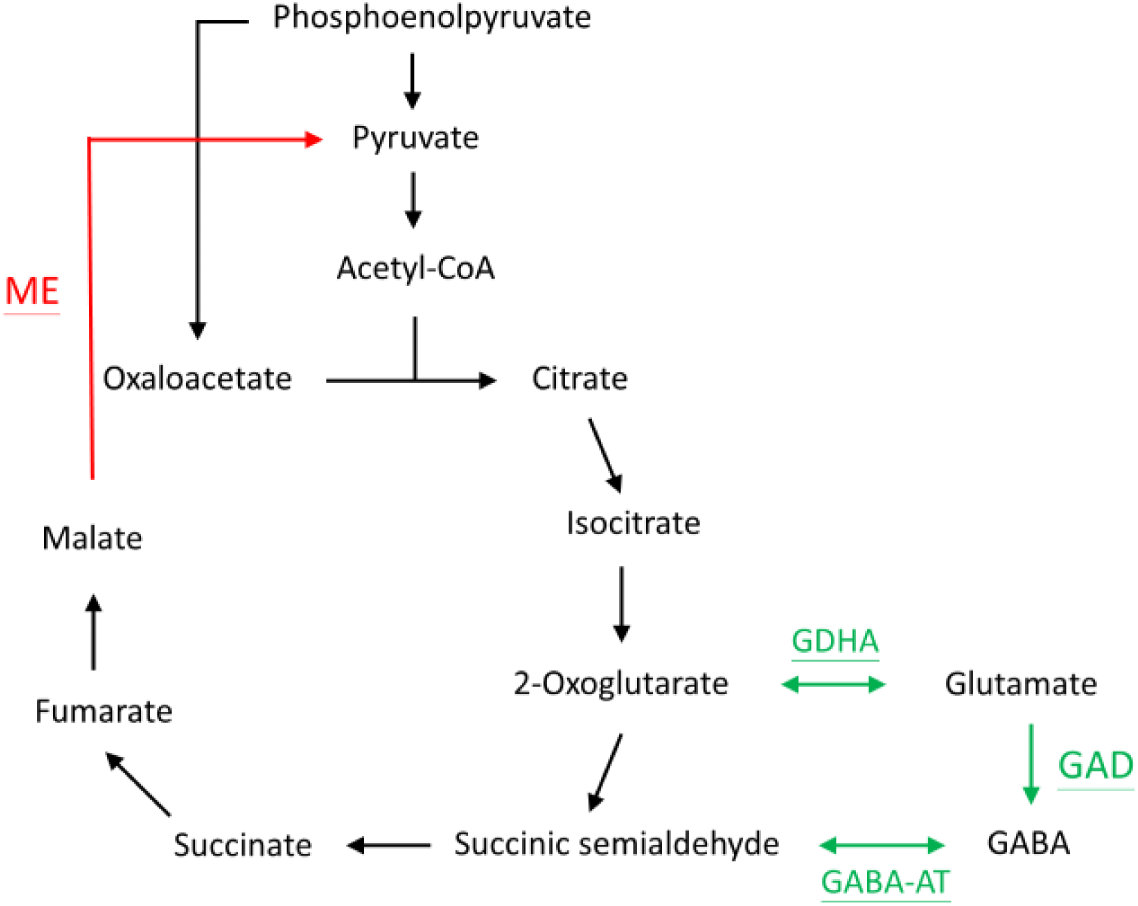
Schematic model of the malic enzyme (ME)-dependent tricarboxylic acid (TCA) cycle and the GABA shunt in DS4 and PCC10605. GAD, glutamate decarboxylase; GDHA, glutamate dehydrogenase; GABA-AT, GABA transaminase. This scheme model is modified from Katayama et al., 2022.

### Cyanobacterial starch metabolism

While most cyanobacteria produce soluble glycogen, several species of unicellular nitrogen-fixing cyanobacteria, such as NBRC 102756 and *Cyanothece* sp. ATCC 51142, accumulated insoluble semi-amylopectin as the storage polysaccharides [11, 57]. DS4 has gene homologs for all three branching-enzyme (BE1, BE2, and BE3: Dongsha4_03540, 03220 and 06100) (Figure 6) found in NBRC 102756 and ATCC 51142 which are required for biosynthesis of semi-amylopectin [57]. By contrast, most cyanobacteria, such as PCC 6803 and PCC 7942, produced soluble glycogen as the major storage polysaccharides and contained only one branching enzyme gene (BE1) in their genomes [58].

### Hydrogen metabolism

DS4 has a gene cluster for homologs of bidirectional NiFe hydrogenase complex (*hoxEFUYH*) [Dongsha4_03740-03725, 03710] and genes encoding for homologs of Hyp proteins (*hypABCDEF*) (Figure 6) involved in the biosynthesis/maturation of these hydrogenases [59]. This NiFe hydrogenase complex may utilize the hydrogen production from nitrogen fixation processes to reduce the soluble electron carriers NADP or ferredoxin.

In addition, DS4 has a distinct gene cluster for homologs of heterodisulfide reductase complex (*hdrABC*) which is consisted of 5 heterodisulfide reductase genes arranged as *hdrB-hdrC-hdrA-hdrB-hdrC* (Dongsha4_08945-08965) (Figure 6). HdrABC plays important functions in the energy metabolism of methanogenic archaea and were found in genomes of few species of cyanobacteria [60]. In most methanogenic archaea, HdrABC formed a complex with methyl-viologen reducing Ni/Fe hydrogenase (MvhAGD) and catalyzed reduction of the mixed disulfide (CoM–S–S–CoB) of the two coenzymes (M and B) and ferredoxin by H_2_ for methane formation [60]. The future study is required to identify reactions and substrates that were catalyzed by HdrABC in DS4.

### Sulfur metabolism

The DS4 genome has a distinct gene cluster for sulfate ABC transport system (*cysPTWA*, Dongsha4_11405-11425) for sulfate uptake which was absent in PCC 10605 (Figure 6). In addition, DS4 and PCC10605 have the full set of genes (adenyl sulfate transferase, adenyl sulfate kinase, phospho-adenosine phosphor-sulfate reductase and sulfite reductase) involved in assimilatory sulfate reduction [61]. Furthermore, DS4 encodes a putative sulfide quinone oxidoreductase (SQR, Dongsha4_00975) which may oxidize sulfide to sulfur for anoxygenic photosynthesis or for detoxification of exogenous sulfide under low-O_2_ conditions [62].

### Thermotolerance genes

The DS4 genome has high number of chaperone genes which may confer their thermos-tolerant properties (Figure 6). For examples, DnaK interacts with its co-chaperones DnaJ and GrpE, and assist folding of denatured proteins under heat and other stress conditions [63]. DS4 and PCC10605 genomes have 7 and 6 copies of chaperone protein DnaK (Hsp70) genes, respectively, versus 3 copies in PCC6803. In addition, DS4 and PCC10605 have 9 copies of chaperone protein DnaJ (Hsp40) genes versus 4 copies in the PCC6803 genome. DS4 and PCC10605 also have 2 copies of chaperone protein GrpE genes versus a single copy in the PCC6803 genome.

In addition, the DS4 genome has the squalene-hopene cyclase (SHC) gene (Dongsha4_04150). The products of SHC, the hopanoids, condense lipid and provide stability of cell membranes under high temperatures and extreme acidity environments due to the rigid ring structures [64].

### Salt and osmotic stress tolerance genes

DS4 exhibits euryhaline (high-salt and osmotic stress tolerant) properties (Figure 1). The basic mechanism of salt acclimation in most cyanobacteria involves the accumulation of compatible solutes and the active extrusion of toxic inorganic ions [65, 66]. DS4 has the glucosylglycerol-phosphate synthase gene (*ggpS*, Dongsha4_01860) for the biosynthesis of the osmolyte glucosylglycerol as well as genes for the metabolism of glycerol-3-phosphate, a precursor of glucosylglycerol [glycerol-3-phosphate dehydrogenase (*glpD*, Dongsha4_00335)]. The accumulation of glucosylglycerol inside cells enabled cyanobacteria to tolerate salinities twice as high as that of seawater (about 6.6% NaCl) [65, 66].

Na^+^ is the main inorganic cation in oceans and in most other saline environments. DS4 has genes for NhaS3 type (Dongsha4_00020, 07485) and NhaP type (Dongsha4_05870, 01470) sodium/proton (or cation/proton) antiporters [67]. In addition, DS4 has genes (Dongsha4_01185-01165, 06870, 06875) for the multi-subunit cation/proton antiporter (Mrp system, composed of ABCDEFG subunits), which may function in osmotic stress tolerance under low salt conditions [68].

Potassium ion (K^+^) transport is also critical for acclimation of cyanobacterial cells to salt and osmotic stresses [67, 69]. DS4 has a distinct gene cluster (*kdpDFABC*) for the high affinity Kdp potassium uptake system (Dongsha4_15640-15655 and 15670) (Figure 6) [69]. In addition, DS4 has the other type of potassium ion transporter system (TrkAH, Dongsha4_03140, 03145, 06635).

Furthermore, DS4 has a predicted gene for the Na-K-Cl cotransporter (NKCC) (Dongsha4_11575) (Figure 6) which may catalyze the coordinated symport of chloride with sodium and potassium ions across the cell membrane in an electrically neutral manner [70].

### Photoprotection

DS4 has four flavodiiron protein homologs (Flv1-4, Dongsha4_14815, 05170, 09665 and 05180) which play important photo-protective roles in photosynthesis of cyanobacteria [71]. Previous studies in PCC 6803 showed that Flv1/Flv3 mediated an oxygen-dependent alternative electron flow under high CO_2_ condition, whereas Flv2/Flv4 was induced and mediated O_2_ photoreduction under CO_2_-limited conditions [71, 72]. Of note, Flv2/Flv4 was absent in the other closely related species (Figure 6). By contrast, these closely related species all contained the catalase/peroxidase HPI gene which might substitute for the function of Flv2/Flv4.

DS4 also has a distinct gene operon (Dongsha4_07235, 07230) encoding for orange carotenoid binding protein (OCP) and fluorescence recovery protein (FRP) which involved in blue-green light induced photo-protection mechanism of cyanobacteria, known as non-photochemical fluorescence quenching [73]. Interestingly, the OCP gene was truncated in PCC10605 (Figure 6).

Superoxide dismutases (SODs) remove superoxide free radicals produced from photosynthetic and respiratory electron transport chains in cyanobacteria. DS4 and PCC10605 have both iron-type (*sodA*) and nickel-type (*sodN*) superoxide dismutase genes (Dongsha4_10025 and 18425; CYAN10605_RS10260 and 11100) to detoxify harmful superoxidemolecules generated in photosynthetic cells. By contrast, PCC7202, HL92 and NIES-3708 have iron-type (*sodA*) only (Figure 6). Most cyanobacterial strains with *sodN* live in saltwater habitats and may have the advantage to grow under iron-limited marine environments [74].

### Novel secondary metabolite genes

Marine cyanobacteria are important resources for pharmaceutically important natural products [75]. We identified the presence of secondary metabolite genes/gene clusters, including nonribosomal peptide (NRP), lantipeptide/bactericide, beta-lactone, and terpenes, in the DS4 genome by using antiSMASH v7.0.0 (see Figure 6).

DS4 has one unique nonribosomal peptide synthetase (NRPS) gene cluster (Dongsha4_00260-00280) which are absent in PCC10605 and related strains (Figure 6). This gene cluster is consisted of 2 amino acid adenylation domain-containing protein genes, as well as genes for phosphopantetheine-binding protein, AMP-binding protein and one hypothetical protein.

In addition, DS4 and PCC 10605 genomes have a novel lantipeptide biosynthesis gene cluster (Dongsha4_03920-03950) (Figure 8) [76]. This gene cluster is consisted of six copies of putative precursor peptide genes and one lantipeptide synthetase gene lanM (Supplementary Figure 4). Precursor peptides contain an N-terminal leader sequence and a C-terminal core peptide. Typical precursor peptides of lantipeptides in cyanobacteria and bacteria are rich in Cys residues which are involved in cyclization of lantipeptides. However, the precursor peptides of DS4 and PCC 10605 are rich in Ser/Thr residues and do not contain any Cys residue (Figure 8). Thus, the products of gene clusters in DS4 and PCC 10605 are expected to produce a novel type of linear D-amino acid containing lantiptides [77].

**Figure 8.**
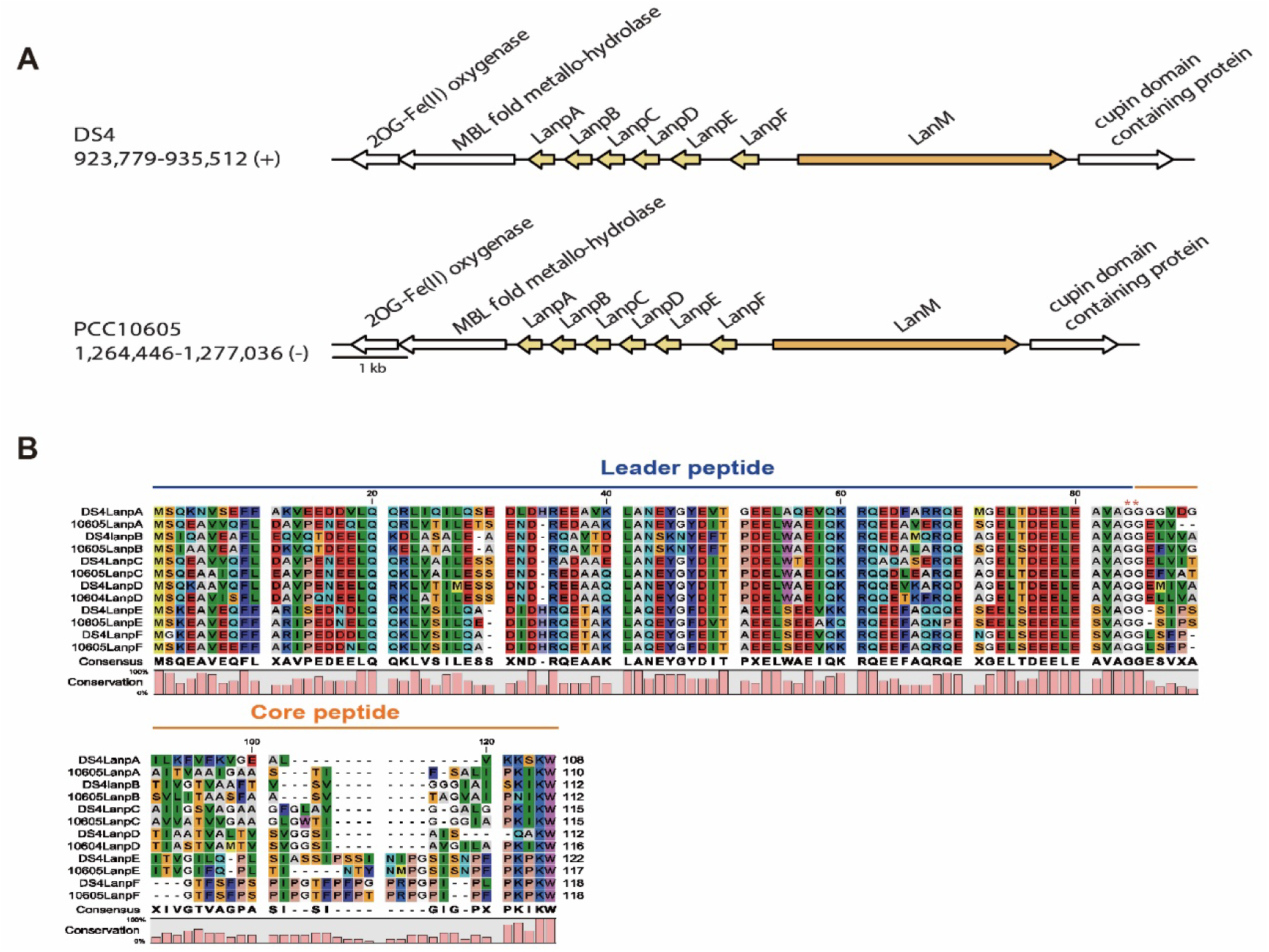
Type 2 lantipeptide synthetase and precursors in DS4 and PCC10605. (A) Synteny plots of lantipeptide biosynthesis gene clusters. Chromosomal locations are labelled below strain name abbreviations on the left. (B) Comparison of amino acid sequences of lantipeptide precursors in DS4 with PCC10605. Precursor peptides contain an N-terminal leader sequence and a C-terminal core peptide. Two conserved glycine residues are labeled with red stars.

Under iron-limiting conditions, many cyanobacteria secrete iron-chelating compounds (siderophores) to sequester the iron essential for growth from the environment [78]. One of the distinct differences between the DS4 and PCC10605 genomes is that PCC10605 has a unique NRPS-independent siderophore synthetase gene cluster that is involved in biosynthesis of the IucA/IucC family siderophores (Figure 6) [78]. In addition, siderophore uptake across the outer membrane in cyanobacteria is usually transported by utilizing TonB-dependent receptors (TBDRs) in the outer membrane [79]. Of note, the PCC 10605 genome has 10 TBDR genes, with none found in the DS4 genome. Furthermore, PCC 10605 has a unique gene cluster for the ABC-type Fe^3+^-siderophore (the hydroxamate type) transport system. Thus, DS4 and PCC 10605 have different types of mechanisms for iron uptake.

## Conclusions

In this work, we reported distinctive genetic features and stress tolerance mechanisms of a thermotolerant and euryhaline strain of unicellular diazotrophic cyanobacterium DS4 isolated from a high-temperature lagoon at Dongsha Island. Unlike related model strains, DS4 has the capacity for nitrogen fixation, as demonstrated by comparative genomics. Additionally, DS4 possesses several distinct genetic features related to special metabolisms, stress tolerant mechanisms, and novel nature product biosynthesis. These findings deepen our understanding of the physiology, ecology, evolution, and stress tolerance mechanisms of cyanobacteria and provide valuable genomic resources for future functional studies and biotechnology applications of heat-tolerant and euryhaline marine cyanobacteria.

### Abbreviations

ANI: average nucleotide identity
BE: branching-enzyme
DS4: Cyanobacterium sp. Dongsha4
FAAL: fatty acyl-AMP ligase
Flv: flavodiiron protein
FRP: Fluorescence recovery protein
GABA: γ--aminobutyric acid
GAD: glutamate decarboxylase
HdrABC: heterodisulfide reductase complex
KGD: α-ketoglutarate dehydrogenase
MDH: malate dehydrogenase
mBG11: modified BG11
ME: malic enzyme
NKCC: Na-K-Cl cotransporter
NRPS: nonribosomal peptide
OCP: orange carotenoid binding protein
PCC10605: *Cyanobacterium aponinum* strain PCC 10605
SEM: Scanning Electron Microscopy
SHC: squalene-hopene cyclase
SOD: Superoxide reductase
SQR: sulfide quinone oxidoreductase
TBDR: TonB-dependent receptor
TCA cycle: tricarboxylic acid cycle
TEM: Transmission Electron Microscopy.

## Declarations

### Ethics approval and consent to participate

Not applicable

### Consent for publication

Not applicable

### Availability of data and materials

The complete genome sequence of *Cyanobacterium* sp. DS4 has been deposited in GenBank/ENA/DDBJ under the accessions CP084098-CP084102. The raw reads have been deposited at the NCBI Sequence Read Archive under the accession PRJNA765482.

## Authors’ Contributions

C-NC, C-HK, and H-AC: conceptualization, funding acquisition, project administration, and supervision. TCY, K-ML, Y-CL, H-YC and Y-FC: investigation, validation, and visualization. C-NC: biological materials. TCY, K-ML, Y-CL, C-NC, C-HK, and H-AC: writing—original draft. All authors: writing—review and editing. All authors contributed to the article and approved the submitted version.

## Acknowledgements

The Illumina and Oxford Nanopore sequencing library preparation services were provided by the Genomic Technology Core (Institute of Plant and Microbial Biology, Academia Sinica). We thank Dr. Wen-Dar Lin in the Bioinformatics Core and Dr. Wann-Neng Jane in the Cell Biology Core (Institute of Plant and Microbial Biology, Academia Sinica) for technical assistance.

