## Supplemental Materials for "Comparative genomic analysis of a novel heat-tolerant and euryhaline strain of unicellular marine cyanobacterium *Cyanobacterium* sp. DS4 from a high-temperature lagoon"

**Supplementary Figure 1.** (A) Determination of *Cyanobacterium* sp. DS4 cell mortality rate using Sytox Green fluorescent dye [1]. These cells were stressed at high temperatures for 3 hours followed by the microscopic imaging using a 100X DIC objective lens. In the right photo, the red fluorescence was chlorophyll emission from the live cells excited by blue light, while the green fluorescence emitted from the dead cells stained by Sytox Green.

(B) Mortality rates of DS4 cells treated in heat stress for 3 hours. The cultures of DS4 cells were under continuous 100 μmol photon $m^{-2}s^{-1}$ white light illuminated from one side of the bottles in the modified B3N medium supplemented with 3% salt. These cells were grown in 1-L transparent glass bottles with 9.5 cm diameters.

(A)

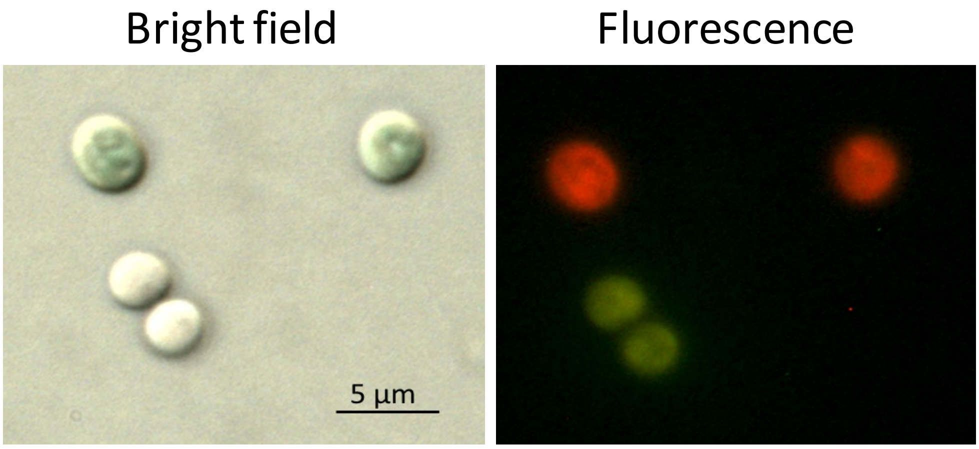


(B)

| Temperature (°C) | Cell mortality (%, mean ± S.E.) | |
|---|---|---|
| | 0 hr | 3 hr |
| 50 | 1.95 ± 0.18 | 1.37 ± 0.30 |
| 60 | 1.93 ± 0.41 | 4.01 ± 0.55 |
| 70 | 0.99 ± 0.14 | 7.05 ± 1.81 |

**Supplementary Figure 2.** (A) Maximum likelihood phylogeny of sentinel D1 homologs based on protein sequences. The locus tag of the DS4 homolog is Dongsha4_01220, GenBank accession numbers are provided for all other strains. Bootstrap support was inferred based on 1,000 replicates, branches with at least 70% support are labeled with the support values. (B) Multiple sequence alignment of photosystem II D1 protein variants. Variations on conserved amino acid ligands to the oxygen-evolving complex in the D1 protein variant psbA4 (sentinel D1 Protein) are labeled with the red stars.

(A)

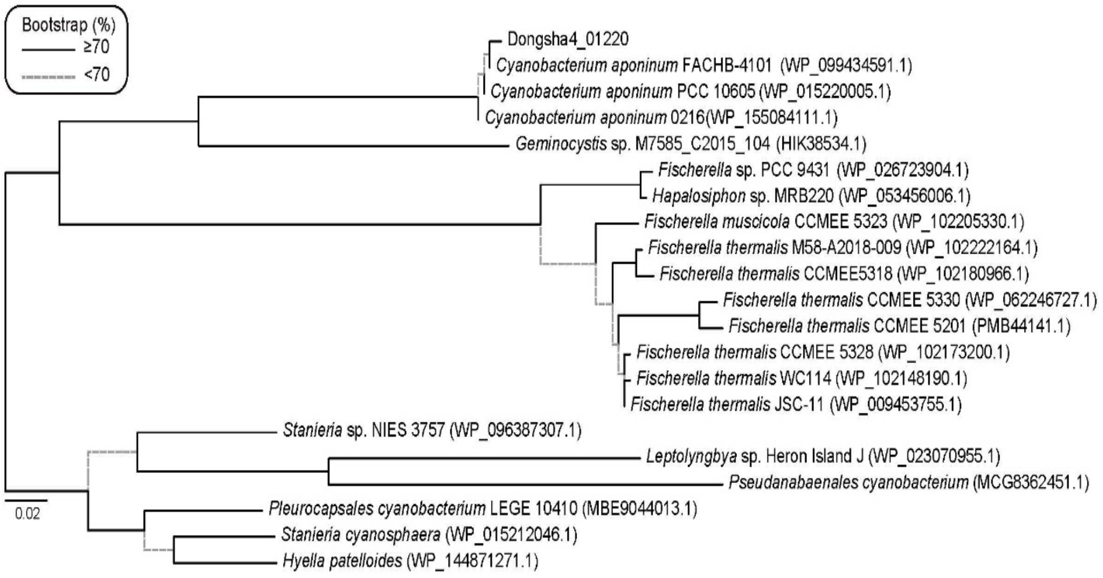


(B)

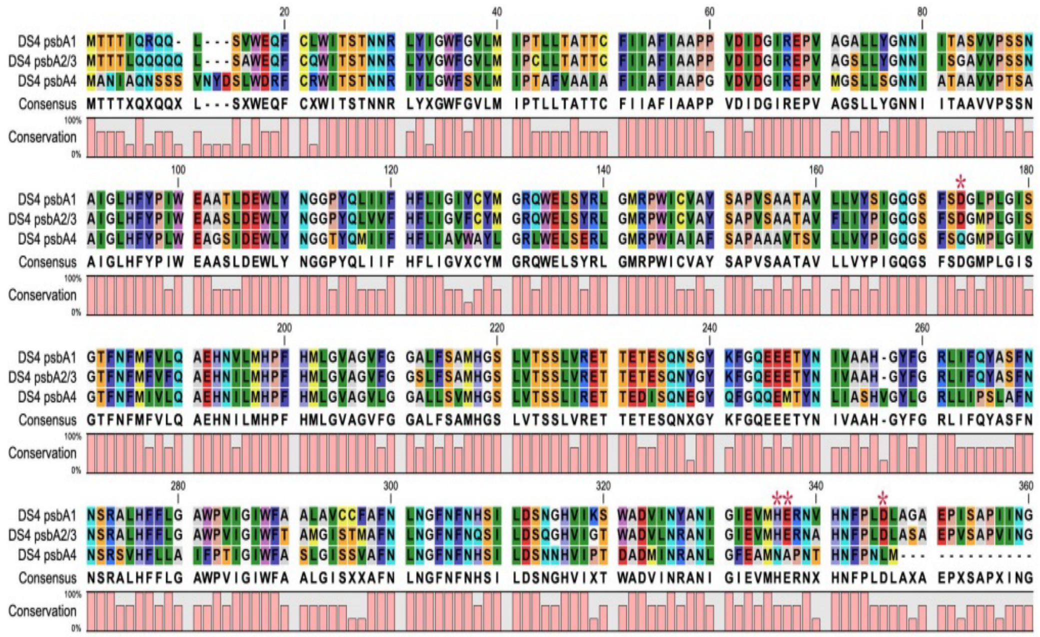

**Supplementary Figure 3.** Maximum likelihood phylogeny of nitrogenase iron protein (NifH) homologs based on protein sequences. The locus tag of the DS4 homolog is Dongsha4_16250, GenBank accession numbers are provided for all other strains. Bootstrap support was inferred based on 1,000 replicates, branches with at least 70% support are labeled with the support values.

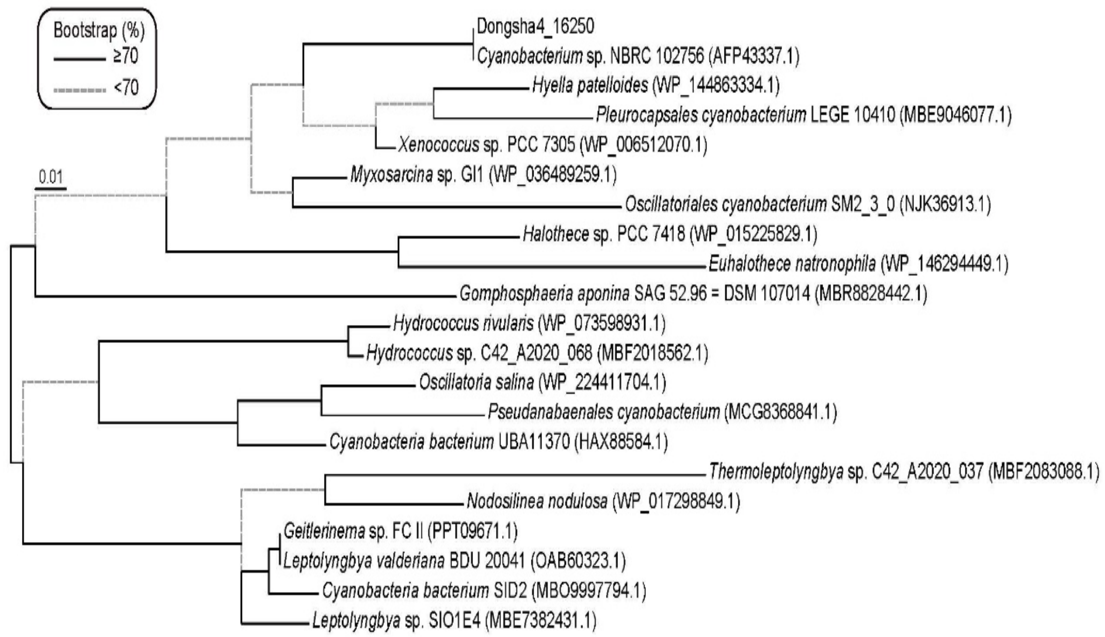

**Supplementary Figure 4.** Maximum likelihood phylogeny of type 2 lantipeptide synthetase LanM family protein homologs based on protein sequences. The locus tag of the DS4 homolog is Dongsha4_03950, GenBank accession numbers are provided for all other strains. Bootstrap support was inferred based on 1,000 replicates, branches with at least 70% support are labeled with the support values.

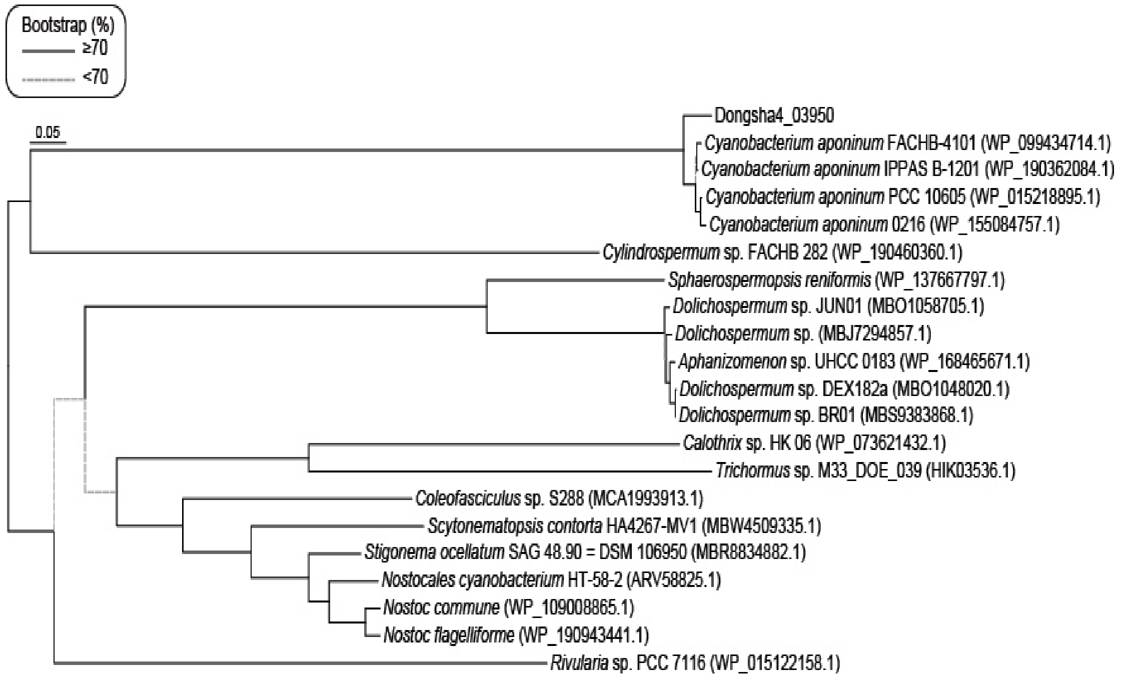
